# Data-driven regularisation lowers the size barrier of cryo-EM structure determination

**DOI:** 10.1101/2023.10.23.563586

**Authors:** Dari Kimanius, Kiarash Jamali, Max E Wilkinson, Sofia Lövestam, Vaithish Velazhahan, Takanori Nakane, Sjors H.W. Scheres

## Abstract

Macromolecular structure determination by electron cryo-microscopy (cryo-EM) is limited by the alignment of noisy images of individual particles. Because smaller particles have weaker signals, alignment errors impose size limitations on its applicability. Here, we explore how image alignment is improved by the application of deep-learning to exploit prior knowledge about biological macromolecular structures that would otherwise be difficult to express mathematically. We train a denoising convolutional neural network on pairs of half-set reconstructions from the electron microscopy data bank (EMDB) and use this denoiser as an alternative to a commonly used smoothness prior. We demonstrate that this approach, which we call Blush regularisation, yields better reconstructions than existing algorithms, in particular for data with low signal-to-noise ratios. The reconstruction of a protein-nucleic acid complex with a molecular weight of 40 kDa, which was previously intractable, illustrates that regularisation through denoising will expand the applicability of cryo-EM structure determination for a wide range of biological macromolecules.

## Introduction

Despite rapid progress in the past decade [1], many biological macromolecules of interest are still too small to allow reliable cryo-EM structure determination. To limit the damage that electrons cause to the biological structures of interest, cryo-EM images are taken under low-dose conditions, which leads to high levels of experimental noise. The noise in the images impedes their alignment, resulting in an ill-posed optimisation problem, where many (noisy or artifactual) reconstructions are equally likely given the data. Nevertheless, the correct solution may still be identified through the incorporation of prior knowledge in the reconstruction process. Most cryo-EM structures are calculated using explicit regularisation of a likelihood function in Fourier space, which assumes cryo-EM reconstructions are smooth in real space [2–4].

Although we know a lot more about the structures of biological macromolecules than that their density varies smoothly, it has been difficult to incorporate richer sources of prior knowledge in the optimisation. Denoising convolutional neural networks provide a mechanism to incorporate complicated prior knowledge into an iterative optimisation process [5]. By training a denoising network on simulated pairs of noisy and ground-truth images, we previously provided a proof-of-principle that prior knowledge about protein structures can be exploited to improve cryo-EM structure determination [6]. However, we also observed problems with overfitting and the hallucination of protein-like features in the resulting reconstructions. Moreover, because experimental cryo-EM structures often comprise regions of well-ordered proteins and nucleic acid domains alongside less structured regions, including for example membrane patches or flexible domains, it was not clear how one would generate ground-truth pairs for experimental cryo-EM data.

Here, we demonstrate how denoising convolutional neural networks, trained and deployed in an application-specific manner, can improve cryo-EM structure determination with experimental data. Similar to the noise2noise approach [7], we trained a denoiser using pairs of noisy reconstructions that are generated as part of standard procedures in cryo-EM structure determination [8]. This approach improves cryo-EM structure determination of a wide variety of biological macromolecules and reduces current size limitations of cryo-EM structure determination.

## Method

## Rationale

The noise2noise framework [7] facilitates the training of a denoising convolutional neural network in the absence of explicit access to ground-truth images. Instead, it relies on pairs of noisy images to extract information about their shared signal. Here, we present an application-specific approach that incorporates this aspect from the noise2noise framework. We trained a denoiser on a set of 422 pairs of noisy half-maps that we downloaded from the electron microscopy data bank (EMDB) [9]. Only entries with reported resolutions higher than 4 Å that had both unfiltered half-maps deposited where selected.

We tailored data augmentation and training of the denoiser to integrate with the iterative expectation-maximisation algorithm for cryo-EM reconstruction. All pairs of half-maps, 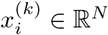, with *k* ∈ {0, 1}, were re-scaled to a uniform voxel size of 1.5 Å, and augmented by generating new pairs 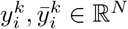:

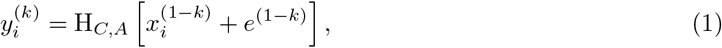

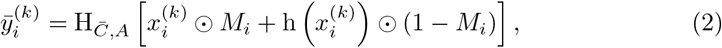

where, *e* ∈R^*N*^ is random coloured noise, *M*_*i*_ ∈[0, 1]^*N*^ is a smooth mask encapsulating the molecules of interest, ⊙represents voxel-wise multiplication, and h(.) is a low-pass filter to 15 Å. H_*C,A*_[.] applies an anisotropic Gaussian filter with covariance matrix *C*, an affine transform *A* that includes rotation and translation, a crop to a cube of 64^3^ voxels, and a voxel-value standardisation. Data augmentation was achieved through random assignments of *C*, 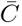, *A, e*, and *r*.

By using a range of resolution cutoffs for *C* and 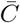, the denoiser explicitly learns to handle maps with varying resolutions. This is needed for its application inside the iterative expectation-maximisation algorithm, which typically starts at relatively low resolutions and gradually progresses to higher resolutions. Although using a lower resolution cut-off for *C* than for 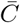 could have produced a network that enhances the resolution of the half-maps, similar to deblurring networks [10], we opted not to do so in order to minimize the risk of hallucinating high-resolution features.

Using different degrees of anisotropy in *C* and 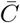, the denoiser learns to deal with the artifacts that arise from non-uniform orientational distributions, while random orientations and affine transformations in *A* lead to invariance with respect to rotations, translations, and intensity scale. By applying h(.) on 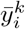, the denoiser learns to enforce fewer features in the solvent region. By filling solvent regions with a 15 Å low-pass filtered version of the map, as opposed to a straightforward voxel-wise multiplication with the mask *M*_*i*_, higher density values in regions with disordered molecules, such as detergent micelles, are maintained.

## Training the denoiser

Our denoiser *f*_*θ*_ consists of a U-net with approximately 13 million trainable parameters *θ* (Figure 1). It is trained using residual learning [11] and with a dropout rate of 50% [12]. Instance normalization [13] is used to handle small mini-batches ℬ, with *b* = 8 samples from the training dataset, during training. We minimise the following loss:

**Fig. 1.**
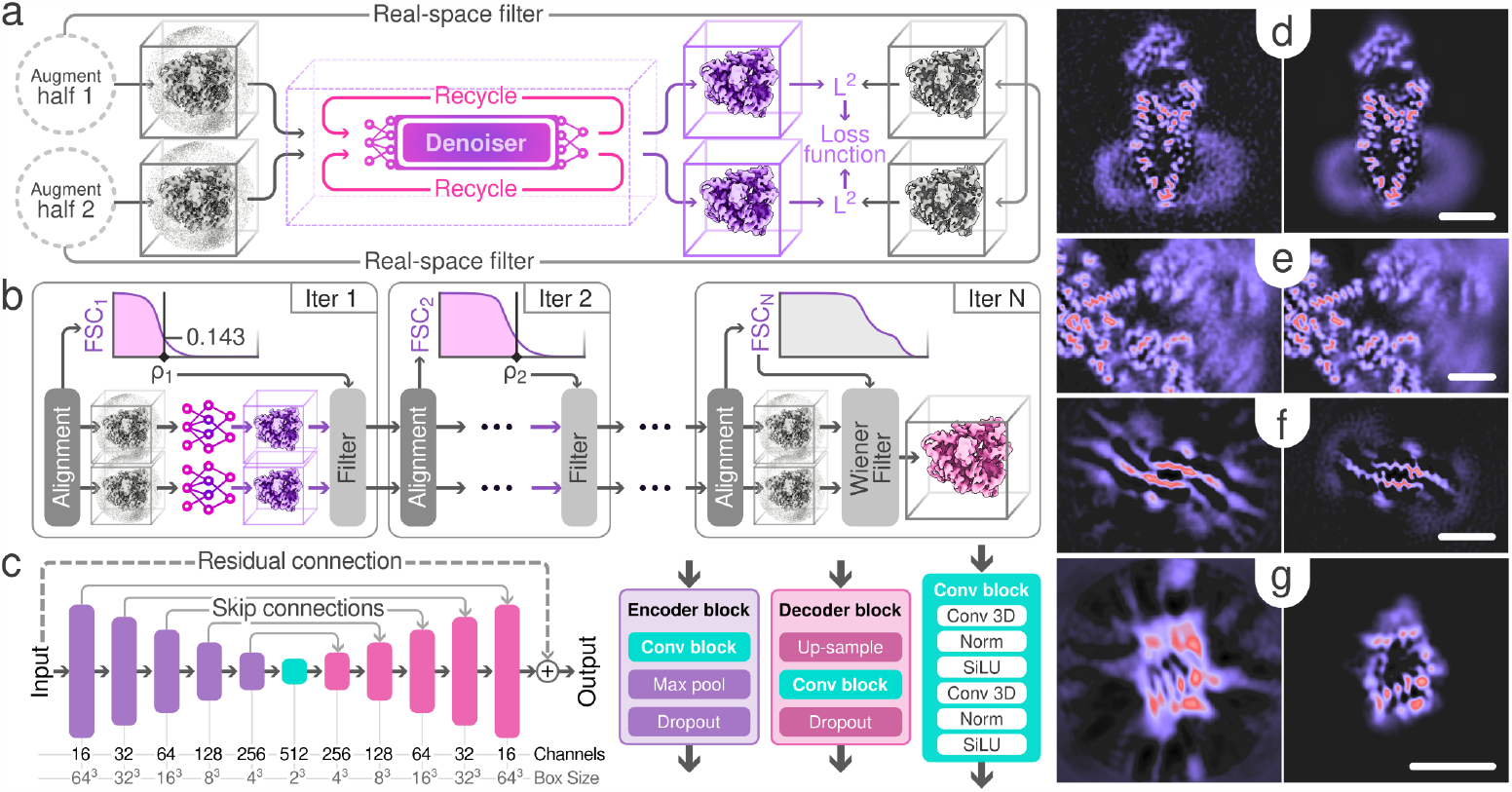
Schematic illustration of Blush and slices of example volumes. **(a)** Training procedure, showing two passes for both half-maps and recycling (R_*r*_) of the denoiser output (in pink). **(b)** Iterative reconstruction with spectral trailing. Each half-map is reconstructed separately. At each iteration, the Fourier Shell Correlation (FSC) is used to estimate a cutoff frequency *ρ*, which is subsequently used to low-pass filter the denoiser output. The final output does not pass through the denoiser but is subject to a Wiener filter, similar to baseline reconstruction. **(c)** Denoiser U-net architecture, consisting of five consecutive encoder blocks, a convolution block, followed by five consecutive decoder blocks. **(d-e)** Slices through maps before (left) and after (right) a single application of the denoiser to the final iteration of the reconstruction for PfCRT **(d)** and the Spliceosome **(e). (f-g)** Showing slices through maps of baseline reconstruction (left) and Blush regularisation (right) of the FIA **(f)** and a 40kDa protein-nucleic acid complex **(g)**. The scale bar indicates 30 Å in all slices.

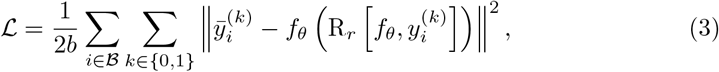

where, R_*r*_[*f*_*θ*_, *y*] returns the output of the denoiser *f*_*θ*_ after recursively calling it *r* ∈ {0, …, 5} times with 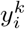 as the initial input. This enables the denoiser to recognize and suppress artifacts brought about by its repeated usage, thereby limiting the amplification of artifacts in the reconstruction that are introduced by the denoiser during subsequent iterations of the expectation-maximisation algorithm [6].

Training for 950,000 steps took 6 days on a single Nvidia A100 GPU.

## Iterative denoising with spectral trailing

Our pre-trained denoiser expresses prior knowledge about cryo-EM reconstructions. Its iterative use within the expectation-maximisation algorithm that underlies cryo-EM structure determination is called *Blush regularisation*. Blush regularisation has been implemented in the open-source software RELION-5. It can be used for 3D classification, multi-body refinement, and 3D auto-refinement jobs, including particles with point-group or helical symmetry.

At every iteration, the denoiser is applied separately to each intermediate half-map reconstruction, where it replaces the 3D Wiener filter that results from regularisation in Fourier space [2, 3]. Although one effect of the denoiser is that it tends to dampen Fourier components at higher spatial frequencies, the amount by which it does so is not well defined. Therefore, we employ a heuristic, here referred to as *spectral trailing*, to prevent overfitting in 3D auto-refinement and multi-body refinement. First, we calculate the Fourier shell correlation (FSC) between two independently refined half-maps before the application of the denoiser and determine the resolution *ρ* where the solvent-corrected FSC drops below 0.143. We then apply the denoiser to both half-maps and subsequently apply a low-pass filter at a resolution that is 2 Fourier shells lower than *ρ*. If *ρ* exceeds the Nyquist frequency of the denoiser, here set to 3 Å, the remaining Fourier shells at higher frequencies are populated with the reconstruction from the standard regularisation in Fourier space. The resulting denoised, low-pass filtered maps are then used as references for alignment in the next iteration. The denoiser is not applied to the output of the final refinement step.

For 3D classification, where data is not separated into independent half-sets, the filtered map from the regularised likelihood approach is used as input for the denoiser. No additional low-pass filtering is applied. In this job type, the denoiser is also applied in the last iteration.

## Results

### Blush improves reconstruction without overfitting

We first tested Blush regularisation on a cryo-EM data set (EMPIAR-10330) [14] of the *Plasmodium falciparum* Chloroquine Resistance Transporter (PfCRT) [15]. This data set has been used as a standard to demonstrate the performance of several approaches to reduce overfitting in cryo-EM refinement [16, 17]. Standard refinement using regularized likelihood optimisation in RELION, which we refer to as the baseline, yields an overall resolution of 3.8 Å for this data set.

Application of Blush regularisation (Figure 2) yielded an overall resolution estimate of 3.4 Å. Spectral trailing was applied with a cut-off at 3.5 Å, beyond which no information from the denoiser was used. Compared to the baseline reconstruction, local resolution improved for most regions of the map, with a corresponding increase in visible side-chain densities. The improvement in resolution as measured by half-map FSC was confirmed by FSCs between both maps and the atomic model that was deposited for this dataset (PDB-ID 6UKJ). Throughout this paper, FSCs between the map and atomic model are calculated using Servalcat [18]. In addition, we also assessed the relative quality of both maps by application of our automated model-building software ModelAngelo [19], which generated a model with 84% completeness in the baseline map and 97% completeness in the Blush map. Model completeness is defined as the percentage of residues that match the reference model with a C*α*-distance of 3 Å or less.

**Fig. 2.**
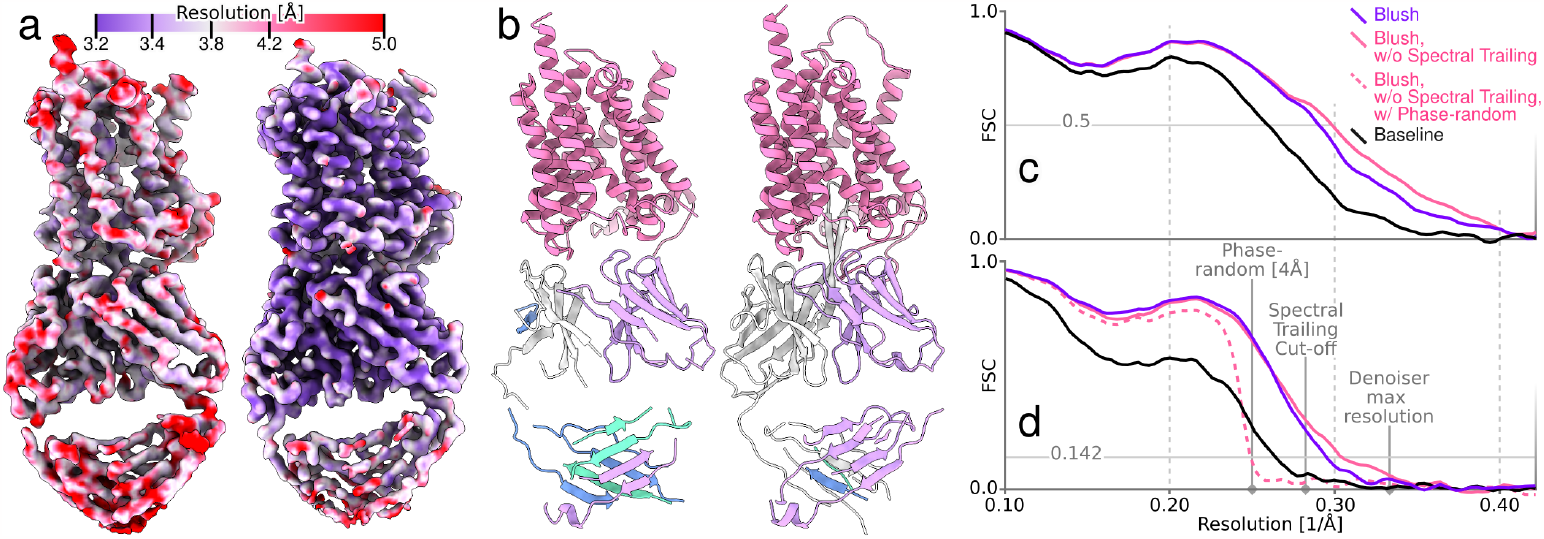
Single-particle reconstruction of the PfCRT dataset. **(a)** Maps coloured by local resolution, comparing baseline (left) and Blush (right). **(b)** Automated atomic modeling by ModelAngelo for the baseline (left) and Blush (right) maps. Colored by chain. **(c)** FSCs between the masked maps and deposited model (ID: 6UKJ). **(d)** Solvent-corrected half-map FSCs. Both plots show FSCs for Blush (purple), Blush without spectral trailing (pink) and baseline (black). Dashed (pink) line show solvent-corrected half-map FSC for Blush without spectral trailing when applied to data with phase randomisation beyond 4 Å resolution.

To assess the potential for overfitting by the denoiser, we also performed a phase-randomisation test [20]. We applied Blush regularisation without spectral trailing for refinement of the PfCRT dataset with phase-randomisation beyond 4 Å. Even though spectral trailing was not used, no overfitting was observed. Switching off spectral trailing led to a marginal improvement in the quality of reconstruction, as quantified by the FSC between the map and the atomic model (Fig 2d). These results indicate that the denoiser can prevent overfitting for this dataset, even without spectral trailing. In the general ase, we still recommend running Blush regularisation with spectral trailing because the gains of switching it off are small and overfitting may be more prominent for other datasets. Consequently, in the following sections, we only present results obtained using spectral trailing.

### Blush expands the applicability of cryo-EM structure determination

We subsequently assessed the broader applicability of Blush regularisation by applying it to four different types of structures and refinement methods.

First, we tested Blush regularisation on a small membrane protein, Ste2, which is a dimeric G-protein coupled receptor (GPCR) [21] (Figure 3, Extended Data Table 1). The full-length monomeric Ste2 has a molecular weight of 47.85 kDa, which includes a long disordered C-terminal tail that comprises 125 amino acids. The total mass of the ordered dimeric Ste2 that contributes to alignment is roughly 67 kDa, most of which lies embedded in a detergent micelle.

**Fig. 3.**
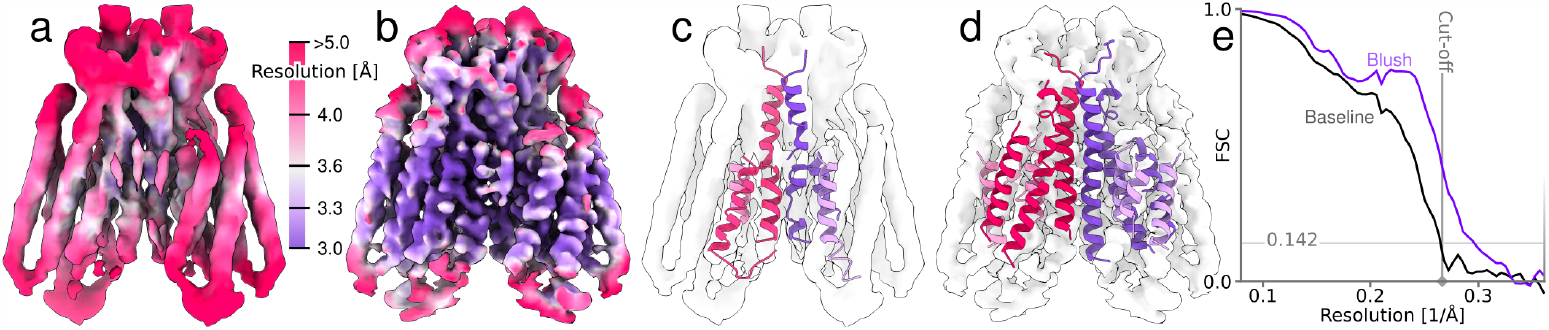
Single-particle reconstruction of the Ste2 dataset. Reconstructions coloured by local resolution, comparing baseline **(a)** and Blush **(b)**. Automated atomic modeling by ModelAngelo, using the baseline **(c)** and Blush **(d)** maps. **(e)** Solvent-corrected half-map FSCs.

The data set was acquired on a similar complex to PDB entry 7QB9 reported in [21], but with different biochemical conditions that impact the stability of the structure. Alignment of images of Ste2 is difficult because not many protein features extend from the smooth detergent micelle. Baseline reconstruction yielded a map with an overall resolution of 3.8 Å, with limited densities for side chains. Application of Blush regularisation led to a structure with an overall resolution of 3.4 Å. Spectral trailing ensured that no information from the denoiser was inserted beyond 3.7 Å resolution. Compared to the baseline reconstruction, the density of the trans-membrane helices is improved. Loops at the top and bottom of the structure are still relatively poorly resolved, probably due to molecular flexibility. In agreement with the visibility of improved side-chain densities and local resolution estimates, the completeness of models built by ModelAngelo in these maps improved from 19% to 43%.

Second, we tested Blush regularisation in multi-body refinement [22]. Multi-body refinement uses partial signal subtraction to align independently moving domains of a larger complex. Reconstructions from subtracted images were not part of the training set of the denoiser. Moreover, signal subtraction reduces the amount of signal in each image, placing stringent limitations on the minimal size of domains that can be aligned in multi-body refinement. We applied Blush in multi-body refinement to a publicly available data set (EMPIAR-10180) of the *Saccharomyces cerevisiae* pre-catalytic spliceosomal B complex [23] (Figure 4). Using four bodies, for the core, the foot, the helicase and the SF3b regions, Blush regularisation improved all domains compared 7 to baseline multi-body refinement, as measured by local resolution, half-map FSCs and FSCs with the reference atomic model (PDB-ID 5NRL). The improvements in resolution were largest in the helicase and SF3b regions, which are the most flexible and thus the hardest to reconstruct. The improvements in resolution were reflected by automated model building in ModelAngelo, which increased model completeness of the entire complex from 32% to 48%. In particular, the model completeness for the SF3b region was improved from 3% to 29%.

**Fig. 4.**
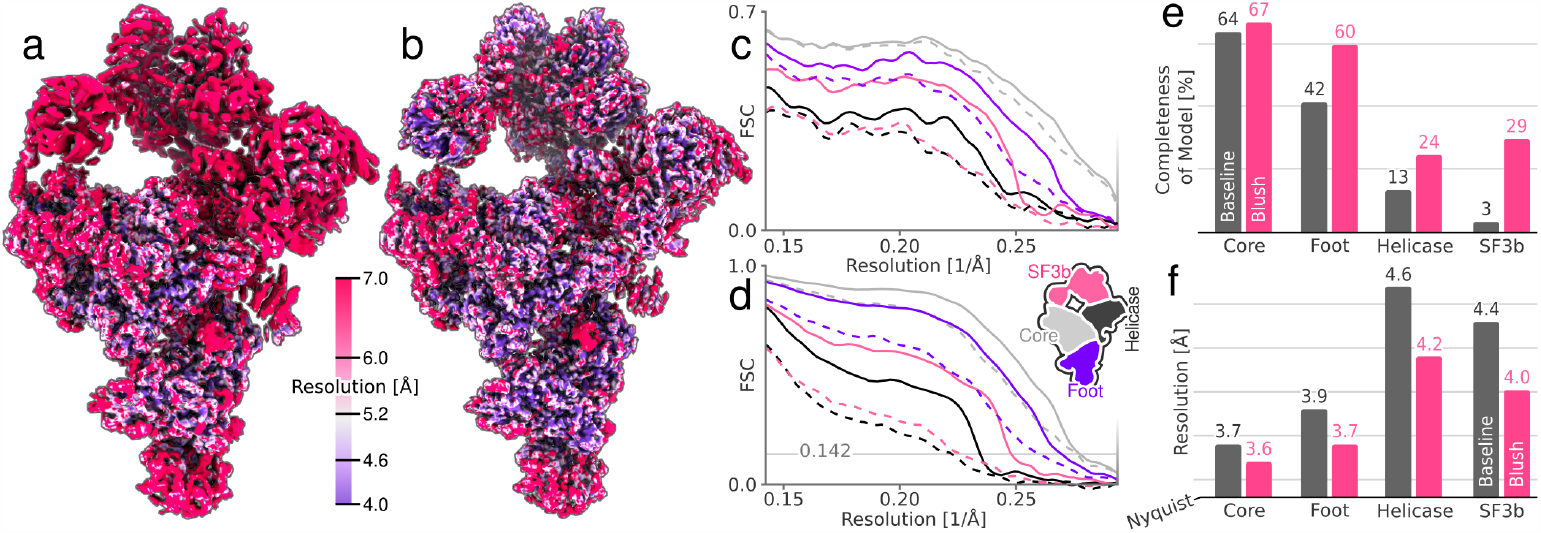
Multibody reconstruction of the spliceosome dataset. Combined maps of the individual bodies, coloured by local resolution, comparing baseline **(a)** and Blush **(b). (c)** FSCs between the masked maps of each body and the corresponding region in the deposited model (ID: 5NRL). **(d)** Solvent-corrected half-map FSCs for the individual bodies. In **(c-d)** dashed and solid lines correspond to baseline and Blush maps, respectively. FSCs are shown for each body: core (grey), foot (black), helicase (purple) and SF3b (red). **(e)** Completeness of atomic models built by ModelAngelo for each body, using baseline (grey) and Blush (red) maps. **(f)** Gold-standard half-map resolutions of each body for baseline (grey) and Blush (red) maps.

**Fig. 5.**
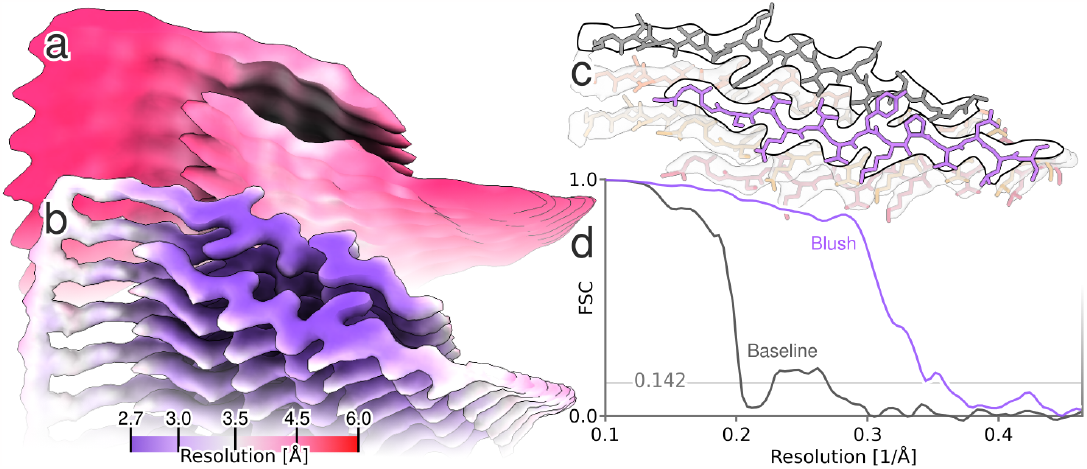
Helical reconstruction of the FIA, coloured by local resolution, for the baseline **(a)** and Blush **(b)** maps. **(c)** Automated atomic modeling by ModelAngelo, comprising tau residues 302-316. **(d)** Solvent-corrected half-map FSCs of the reconstructed maps.

Third, we assessed the performance of Blush regularisation for a biological assembly that differed from the types of structures the denoiser was trained on: the first intermediate amyloid (FIA) that forms during the *in vitro* assembly of recombinant tau (residues 297-391) [24]. This data set is also publicly available (EMPIAR-11720). Unlike any of the structures in the training set, the FIA has helical symmetry. It is an amyloid filament, with parallel *β*-strands repeating every 4.7 Å in the direction of the helical axis. Besides deviating from the types of structures in the training set, the FIA is also one of the smallest amyloid structures solved to date, with only 15 ordered residues in each of two opposing *β*-sheets. Baseline helical refinement yielded a 5.0 Å map, in which the density for *β*-strands along the helical axis was not separated, and no atomic model could be built. Blush regularisation improved the resolution to 2.8 Å, and ModelAngelo built all 15 ordered residues in the resulting map.

Fourth, we applied Blush to a protein-nucleic acid complex with a combined molecular weight of 40 kDa (Figure 6; Extended Data Table 1). Using multiple different classification and refinement strategies in baseline RELION and CryoSPARC, we were unable to obtain a reliable reconstruction (results not shown). Even though an initial model generated using the standard VDAM algorithm in RELION [25] suffered from anisotropy, a first 3D classification with Blush regularisation resulted in one class with recognizable protein features. Refinement of the corresponding class yielded a better initial model for a second 3D classification, from which a single class was selected for subsequent CTF refinement [26] and particle polishing [26]. A 3D classification was performed without alignment, followed by a final 3D refinement. All 3D classifications with alignment and 3D refinements used Blush regularisation. The final map achieved a resolution of 2.5 Å, with ModelAngelo successfully building 97% of the protein sequence, and 33 out of 42 nucleotides.

**Fig. 6.**
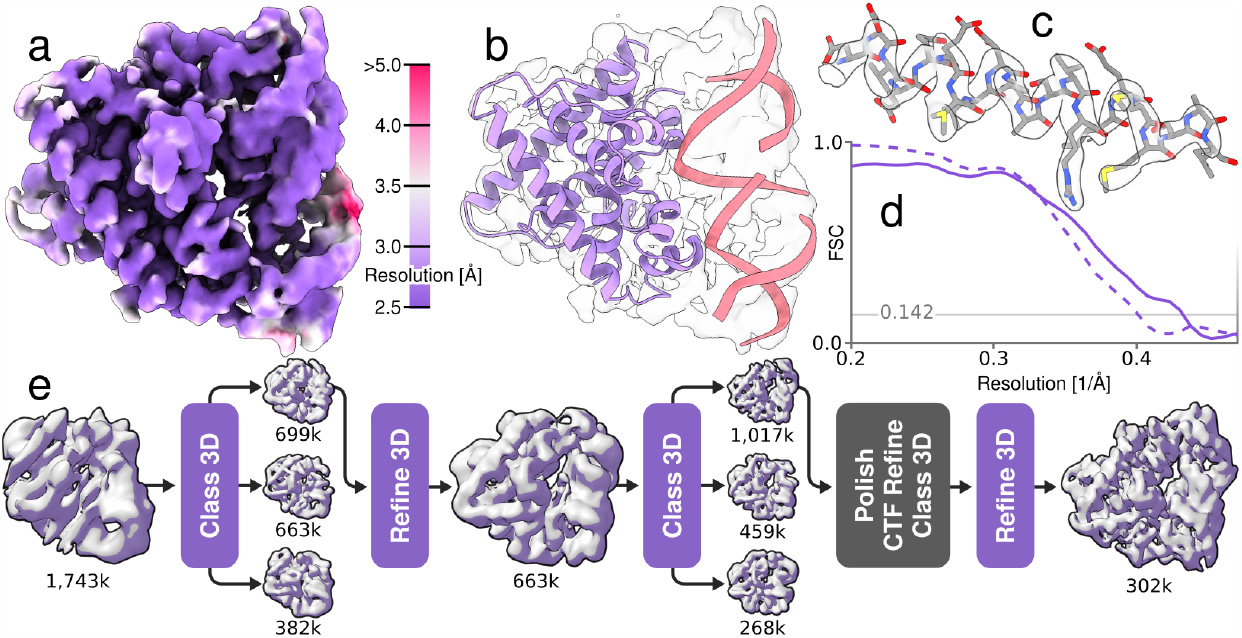
Single-particle reconstruction of the protein-nucleic acid complex with a molecular weight of 40 kDa. **(a)** Blush regularisation local resolution. **(b)** Automated atomic model assignment with ModelAngelo. **(c)** Detailed view of an alpha-helical segment in the reconstructed map and refined atomic model. **(d)** Solid line shows FSC to a reference atomic model and the dashed line shows half-map FSC. Solvent-corrected resolution is 2.8 Å using a spectral trailing cut-off at 3.0 Å. **(e)** Processing pipeline from initial model to final reconstruction. Numbers indicate the number of particles assigned to each map. Purple squares indicate reconstructions using Blush regularization.

## Discussion

Our results demonstrate that denoising convolutional neural networks can be used as a source of extended prior information in the reconstruction of experimental cryo-EM images. A previous approach that used noise2noise, implemented in the M software [27], exclusively trained a denoising network on half-maps from the dataset to which it was also applied. As a result, unlike our approach, it did not express general prior knowledge about cryo-EM reconstructions. When we tried to express prior knowledge about protein structures by training a denoiser on pairs of noisy and ground-truth maps that were calculated from atomic models, we observed problems with overfitting and hallucinations [6]. Similar problems may also explain why the application of the DeepEMhancer neural network [28] inside the iterative reconstruction algorithm of RELION had to be restricted to only a few iterations at the end of refinement [29]. The approach described in this paper reduces the risk of hallucinating protein-like features in the reconstruction by using a neural network that is trained on experimental cryo-EM half-maps only, i.e. without using atomic models or the geometrical restraints that are used to describe them.

Instead of forcing the map to resemble densities derived from atomic models, our denoiser is trained to introduce more subtle modifications to cryo-EM maps, such as smoothing out density in solvent regions or within detergent micelles. The network also removes artifacts that are commonly encountered in difficult cryo-EM refinements, e.g. anisotropic densities that result from uneven angular distributions, or radially extending, streaky features that are often observed in overfitted maps (Figs 1f-g). Our findings illustrate that, even though the effect of a single application of the denoiser is relatively small, its cumulative impact over multiple iterations enhances the performance of cryo-EM structure determination across a diverse range of test cases. As machine-learning methods get better at extracting knowledge from large datasets, it may be tempting to exploit more knowledge about the structures of biological macromolecules in the reconstruction process. However, doing so may ultimately also take away one of the most powerful ways of assessing whether a reconstruction is correct: the presence of expected features in the map. We thus anticipate that the cryo-EM community will continue to explore the question of how much prior knowledge should inform the reconstruction process and how much should be kept aside for validation.

Within the framework of Blush regularisation, the denoiser replaces the exponential prior that traditionally constrains the power of Fourier-space components in the baseline algorithm. As a result, the FSC between independently refined subsets no longer influences the weighting of Fourier components in intermediate reconstructions. Instead, this FSC is used to determine a resolution cutoff *ρ*, beyond which the Fourier components of the two denoised half-maps are set to zero. Because Fourier components near the resolution estimate of the final map will not have been affected by the denoiser, protein-like features at these frequencies cannot be the result of hallucination or overfitting by the denoiser. Future investigations will explore alternative approaches that do not rely on heuristics, like choosing a specific *ρ*, which arises from the observation that the dampening effect of the network is not well defined in Fourier space. Future adaptation of the VDAM algorithm [25] will also allow the use of Blush regularisation for initial model generation.

In all our tests, Blush regularisation outperformed or equaled the baseline implementation in RELION, with the differences being largest for cases where the baseline approach tends to overfit the data. Consequently, Blush regularisation will be most useful for refinements of data sets with low signal-to-noise ratios, like those of small complexes or complexes embedded in thick ice layers, multi-body refinements involving relatively small bodies, and refinements of maps exhibiting pronounced variations in local resolution. For example, Blush regularisation allowed reconstruction of an amyloid with only 30 residues in its ordered core and a protein-nucleic acid complex with a molecular weight of 40 kDa. Although nucleic acids result in higher signal-to-noise ratios than proteins, 40 kDa approaches predictions of the minimum size protein that is amenable to cryo-EM structure determination [30, 31]. These results illustrate that the use of prior knowledge through denoising convolutional neural networks expands the applicability of cryo-EM structure determination.

## Author contributions

DK designed and implemented Blush regularization, ran most experiments and analysed the results. KJ contributed to data preprocessing. MEW contributed and analysed the dataset of the 40kDa protein-nucleic acid complex, and contributed to analysis of the PfCRT dataset. SL contributed the FIA dataset. VV contributed the Ste2 dataset. SHWS supervised the project and contributed to RELION integration. All authors contributed to the writing of the manuscript.

## Acknowledgements

We thank Johannes Schwab, Keitaro Yamashita, Carola-Bibiane Schönlieb, and Ozan Öktem for helpful discussions; Jake Grimmett, Toby Darling, and Ivan Clayson for help with high-performance computing; the EM facility of the Medical Research Council Laboratory of Molecular Biology for support with cryo-EM; and Ed Brignole and Chris Borsa for the smooth running of the MIT.nano cryo-EM facility, established in part with financial support from the Arnold and Mabel Beckman Foundation. MEW is grateful to Feng Zhang for funding support. TN is a member of the JEOL YOKOGUSHI Research Alliance Laboratories. This work was supported by the Medical Research Council as part of the United Kingdom Research and Innovation (MC UP A025 1013 to SHWS); the European Union’s Horizon 2020 research and innovation programme (under grant agreement no. 895412 to DK); a Helen Hay Whitney Foundation Postdoctoral Fellowship (to MEW); the Howard Hughes Medical Institute (to MEW) and a Research Fellowship at Gonville and Caius College of Cambridge University (to VV). For the purpose of open access, the MRC Laboratory of Molecular Biology has applied a CC BY public copyright licence to any Author Accepted Manuscript version arising.

## Code availability

Blush regularisation has been implemented in the open-source software RELION-5, which is distributed for free under the GPLv2 license and can be downloaded from https://github.com/3dem/relion.

## Extended Data

**Extended Data Table 1.** Details on cryo-EM data collection and processing of the Ste2 and 40 kDa complex datasets.

|  | **Ste2** | **40 kDa complex** |
| --- | --- | --- |
| **Data collection** |  |  |
| Microscope | Thermo Scientific Krios G3i | Thermo Scientific Krios G3i |
| Camera | Falcon 4i | Gatan K3 |
| Magnification | 96,000x | 165,000x |
| Voltage (kV) | 300 | 300 |
| Pixel size (Å) | 0.824 | 0.531 |
| Total electron exposure ($e^-/\text{Å}^2$) | 56 | 72 |
| Exposure rate ($e^-/\text{pixel}/\text{sec}$) | 7.9 | 15 |
| Defocus range ($\mu\text{m}$) | 0.8 - 2.4 | 0.8 - 1.5 |
| Automation software | EPU | EPU |
| Energy filter slit width | N/A | 20 eV |
| Micrographs collected (no.) | 13,254 | 11,232 |
| **3D reconstruction** |  |  |
| Number of frames for motion correction | 70 | 40 |
| Total extracted particles (no.) | 2,608,181 | 6,449,613 |
| Final particle images (no.) | 147,797 | 301,833 |
| Symmetry imposed | C2 | C1 |
| Resolution (FSC 0.143; unmasked/masked) | 3.9 / 3.4 | 2.8 / 2.6 |
| Sharpening B-factor ($\text{Å}^2$) | -160 | -90 |

